# *ythdf2(ch200)* and its role in development of the early zebrafish embryo

**DOI:** 10.1101/2022.12.02.516060

**Authors:** Alana V. Beadell

## Abstract

We appreciate the well-presented data and focus on mechanism in the paper titled, “Ythdf m^6^A Readers Function Redundantly during Zebrafish Development” by Kontur *et al*. December 29, 2020^1^ [DOI: https://doi.org/10.1016/j.celrep.2020.108598]. However, we would like to suggest several alternative conclusions regarding the role of Ythdf2 in maternal RNA clearance and the phenotypic consequences of the *ythdf2(ch200)* mutation as described in Zhao *et al*. 2017, “m^6^A-dependent maternal mRNA clearance facilitates zebrafish maternal-to-zygotic transition”^2^ [DOI: https://doi.org/10.1038/nature21355], along with caveats regarding the interpretation of Ythdf2’s roles in mRNA metabolism in the early embryo.

## Methods

### Zebrafish maintenance

Heterozygous mutant *ythdf2* fish in the *AB background were custom generated by ZGeneBio (Taiwan) via TALEN mutagenesis as described in Zhao et al. 2017^2^. Purchased heterozygous males were outcrossed to in-house wild-type *AB stock fish for three generations before being in-crossed to produced homozygous *ythdf2* mutants for study. PCR primers used for genotyping were as follows: ythdf2F: (5’)GACAAATCTGCCACCACCTC(3’) and ythdf2R: (5’)CCACTGCCATTCACACTGAA(3’). Genotyping PCR on purified genomic DNA was performed with Phusion High-Fidelity DNA Polymerase (NEB). PCR products were checked by agarose gel for single band amplification of the appropriate size and then subjected to Sanger sequencing for definitive identification of the 8 bp *ch200* DNA deletion in *ythdf2* exon 4. All embryos studied were obtained from natural crosses, raised under standard conditions (*e.g*., at 28.5°C in E3 medium), and staged according to Kimmel *et al*. 1995^3^. Fish were maintained in accordance with AAALAC research guidelines under a protocol approved by the University of Chicago Institutional Animal Care & Use Committee.

### *In situ* hybridization

Antisense digoxigenin-labelled RNA probes were generated by *in vitro* transcription using the MEGAscript T7 Transcription Kit (Invitrogen) along with DIG RNA Labeling Mix (Roche) and were purified via phenol-chloroform extraction/isopropanol precipitation. *In situ* hybridizations were performed essentially as described in Thisse and Thisse (2014)^4^. All assays were repeated at least once using different biological replicates in separate experiments.

### Microscopy

All images were observed using a Leica MZFLIII microscope and captured with a Nikon D5000 digital camera using Camera Control Pro (Nikon) software. For bright field imaging of live embryos, image saturation values were adjusted identically for all images. No other image processing was performed.

## Results and Discussion

### The blastula-stage developmental delay observed in maternal *ythdf2(ch200)* embryos is not due to genetic background

Kontur *et al*. state that loss of Ythdf2 does not delay early zebrafish development. By in-crossing maternal-zygotic homozygous mutants and comparing their offspring to those produced by an in-cross of “background-matched” fish that house only wildtype Ythdf2 alleles, Kontur *et al*. were unable to observe any phenotypic developmental delay using either the *ythdf2(ch200)* allele (8 base pair deletion; Zhao *et al.)* or a 223 base pair *ythdf2* deletion allele they generated themselves. Both deletions occur in exon 4 of *ythdf2*, the largest exon of this gene and the exon that codes for Ythdf2’s RNA binding domain. Yet, Kontur *et al*. found that when the progeny of in-crossed maternal-zygotic homozygous *ythdf2* mutants were instead compared to progeny of a mating between two different wild-type zebrafish strains, (*AB and Tübingen), they were able to observe a developmental delay in embryogenesis. Along with failure to obtain high-throughput sequencing data showing substantial differences in steadystate mRNA levels between progeny of in-crossed *ythdf2* maternal-zygotic mutants and “background-matched” siblings, Kontur *et al*. conclude that the delay in developmental timing observed in Zhao *et al*. just after the mid-blastula transition (MBT), as well as the substantially different mRNA profiles we obtained at different developmental time points by high-throughput sequencing, must not be due to the *ch200* allele itself, but instead to some natural difference in developmental timing between the genetic background into which the *ch200* allele was originally created and/or crossed and the wild-type “background strain” to which it was compared.

However, this hypothesis is not likely to explain our findings in Zhao *et al*. for a number of reasons. First, as is customary for all generated mutations, and before any observation, we outcrossed the *ythdf2(ch200)* allele (which was created in the *AB background by an outside laboratory) three times to our own *AB wild-type background strain, including at least one time using males as the *ch200* carrier in order to guard against unintended mitochondrial DNA mutations hitchhiking along with *ch200*. This practice not only helps to separate unlinked, secondary mutations from the mutation of interest, but also helps guard against genetic background differences that might confound analyses by homogenizing the initial background of mutant fish to the control fish against which they will be compared. All of our in-house outcrossings of *ch200* were performed using the same population of *AB fish that ultimately served as our controls. We maintained relatively large breeding populations of wild-type and heterozygous and homozygous *ch200* mutant fish (*i.e*., 10 homozygous *ch200* females, ~25 heterozygous *ch200/+* females, and ~50 wild-type *AB females). We did not rely on set breeding pairs, but instead freely mated wild-type and homozygous female mutants to either wild-type or homozygous *ch200* males. Very often, we used mutant embryos from multiple mating pairs in any single experiment, further ensuring that low-frequency alleles at loci other than *ythdf2* were unlikely to influence the blastula-stage developmental delay and molecular phenotypes we observed. Moreover, “developmental delay” is not a mutant phenotype that we have observed previously in our laboratory, although we have studied numerous mutations generated both within and outside of our lab and both unrelated and related to m^6^A metabolism.

Most importantly, direct evidence for the lack of salient genetic background effects in our prior study of Ythdf2 comes from comparing the developmental timing of embryos produced by our control *AB fish versus *m(+/+) z(+/-) ythdf2(ch200)* embryos, which, as described above, had been crossed into our *AB background. As shown in **Figure 1**, we find no differences in early developmental timing between these two genotypes. By contrast, Kontur *et al*. observed differences in the developmental timing between their control *AB-Tübingen hybrid fish progeny and the genetic background of *ch200*. We have not independently tried to confirm this finding, but we emphasize that the Tübingen wild-type zebrafish strain played no part in our experimental design, and we have no reason to consider that Tübingen-specific wild-type alleles are a part of our observations.

**Figure 1.**
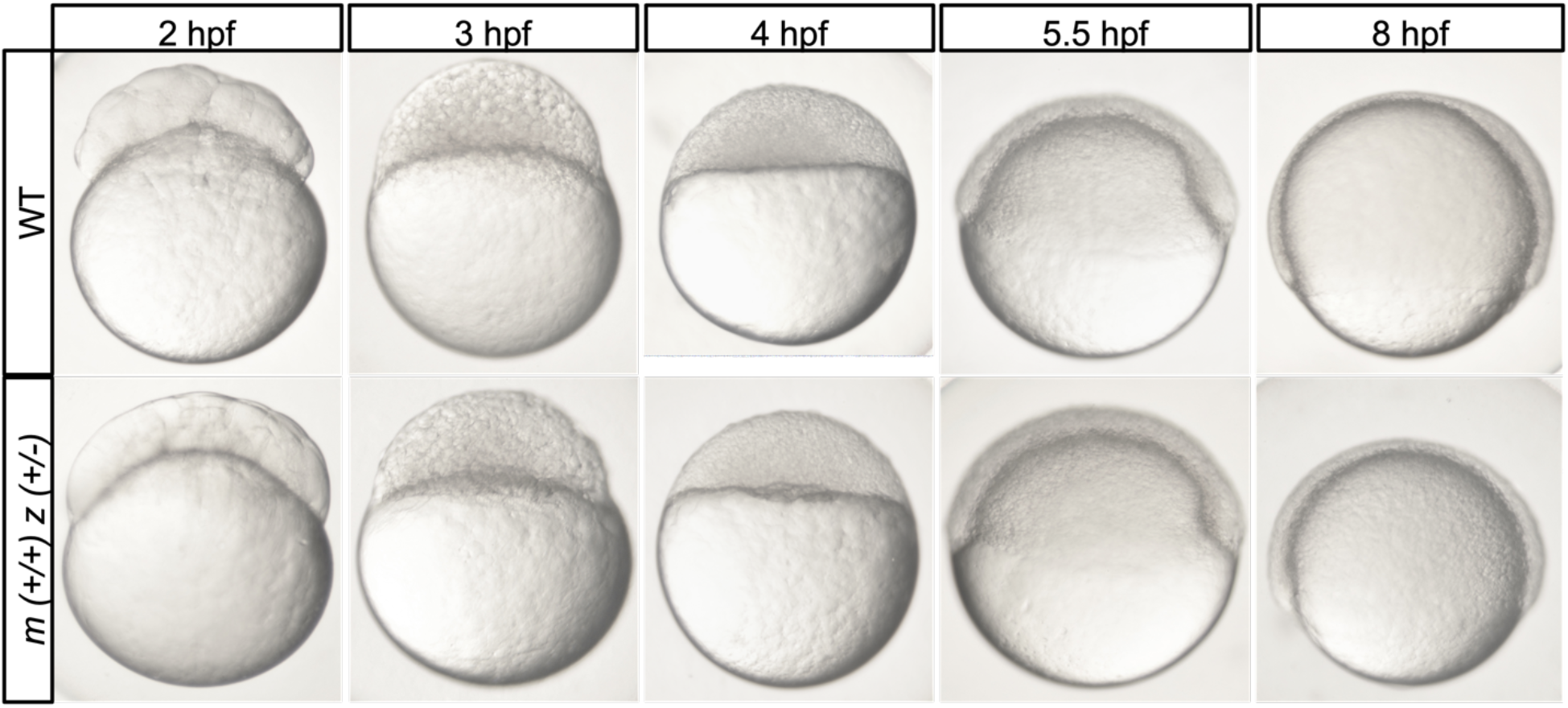
The blastula-stage developmental delay observed in maternal ythdf2(ch200) embryos is not due to genetic background. Performed at the same time as the results published in Zhao *et al*. 2017^*2*^, direct comparison of embryos produced from control wild-type fish (*AB) and from mating *AB females to ch200 heterozygous males (originally generated in the *AB background and subsequently outcrossed to our *AB stock) produces no differences in developmental timing at the stages shown (or later).

To summarize these points, during the course of our work we observed no phenotypic differences between our *AB control embryos and *m(+/+) z(+/-) ch200* embryos (in the *AB background), but we did observe differences between our *AB control embryos and *m(-/-) z(+/-)* or *m(-/-) z(-/-) ch200* embryos (in the *AB background). Together, these comparisons provide a proper control for “genetic background” and, along with the molecular evidence of cell-cycle rescue provided in Extended Data Figure 2 of Zhao *et al*., argue that *ch200* is responsible for the phenotypic and molecular sequencing results that we observed in that work. **Figure 2** provides additional images of the maternal *ch200* developmental delay morphological phenotype in an assay performed two years after the submission of Zhao *et al*. 2017.

**Figure 2.**
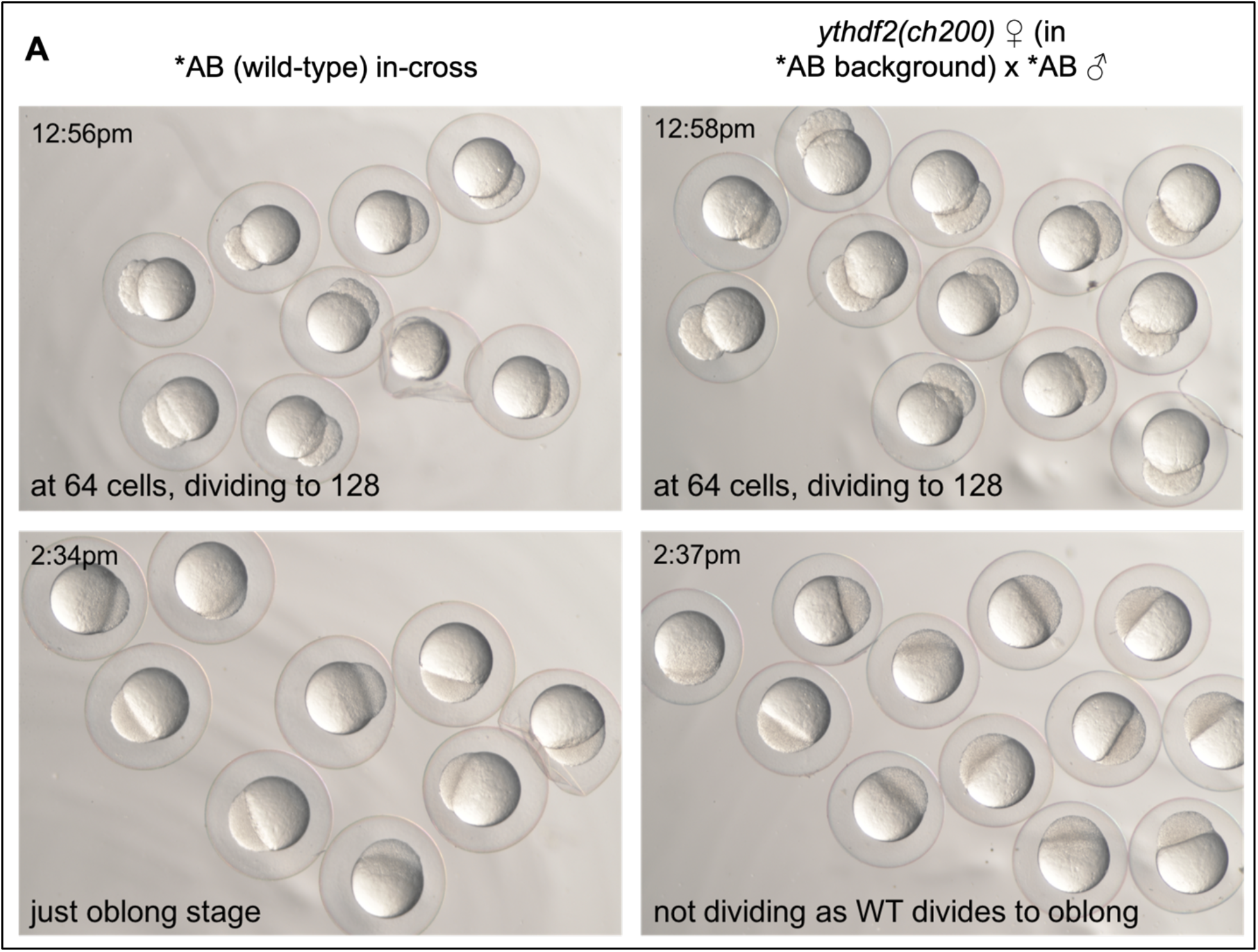
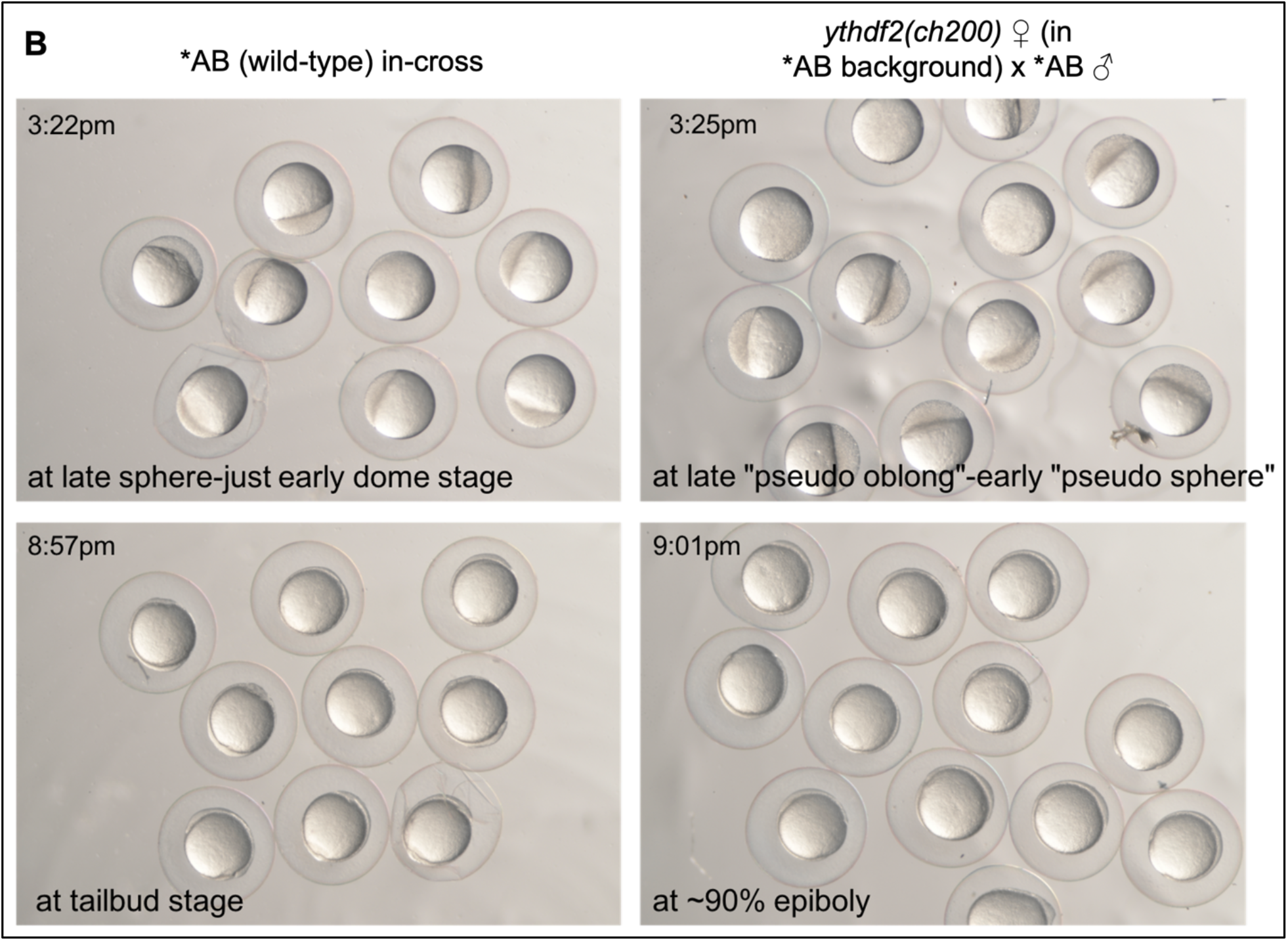
The morphological developmental delay of maternal ythdf2(ch200) embryos is recapitulated in a phenotyping assay performed two years later and in a subsequent generation of fish from Zhao et al. (2017)^2^. (A) Wild-type (*AB) and maternal (-/-) zygotic (+/-) *ythdf2(ch200)* embryos (in the *AB background) photographed at 64 cells (cleavage-stage embryos, before the MBT) and again after MBT when wild-type embryos have just divided to reach the oblong stage, but mutant embryos at the same time have aberrantly paused during cell cycle 12 of embryogenesis; (B) the same embryos as in (A), photographed at the resumption of cell division in mutants (at the same time that wild-type embryos are one stage ahead of mutants, now dividing to dome stage) and again when wild-type embryos have reached the nominal end of gastrulation (the tailbud stage), but mutant embryos are only at 80-90% epiboly.

### If maternally homozygous *ythdf2(ch200)* mutants produce progeny with delayed developmental timing just after MBT, why was this not observed in a subsequent study?

We suggest that to observe the developmental delay phenotype in embryos produced by *ythdf2(ch200)* mothers is not trivial, as cells at the relevant developmental stages (zebrafish high stage through sphere stage) are smaller compared to earlier blastomeres, the cell cycles are relatively short (~20 minutes), and cell division has become a bit asynchronous versus earlier cleavage and blastula stages. Observation of the maternal *ch200* delay mutant phenotype requires that embryos be precisely staged (*i.e*., within 2 minutes of one another in developmental time) many cell divisions in advance of the MBT and that embryos be continuously monitored under identical environmental conditions.

We emphasize that maternal *ch200* blastula-stage embryos skip only a single cell division, such that they remain morphologically static for a total of ~45 minutes (at 29 +/- 1.5°C) during the high stage of embryological development (12^th^ cell cycle), while wild-type embryos divide once during that time interval to proceed from high to oblong stage. Importantly, during the maternal *ch200-* mediated developmental delay, mutant cells *en masse*, but perhaps not every cell, miss only this single cell division. During the course of our observations in Zhao *et al*., we paid careful attention to the observable aspects of different mitotic stages (*e.g*., disappearance of the nuclear envelope) and to the observable reduction of cell size that follows each mitosis at these developmental time points. We found that “cell settling” occurs in the posterior portion of the blastula (nearer to the blastula-yolk cell interface) as mutant embryos near the division to the “pseudo-oblong” and “pseudo-sphere” stages, such that in overall size and shape, mutant embryos may look “oblong” or “sphere” at first glance. **Figure 3** provides a direct comparison between phenotypically wild type-looking and developmentally delayed embryos just after MBT. Accidental mistiming of embryo development or misidentification of developmental stage will render comparative sequencing or other embryo harvesting-based assays performed at those times and stages ineffectual.

**Figure 3.**
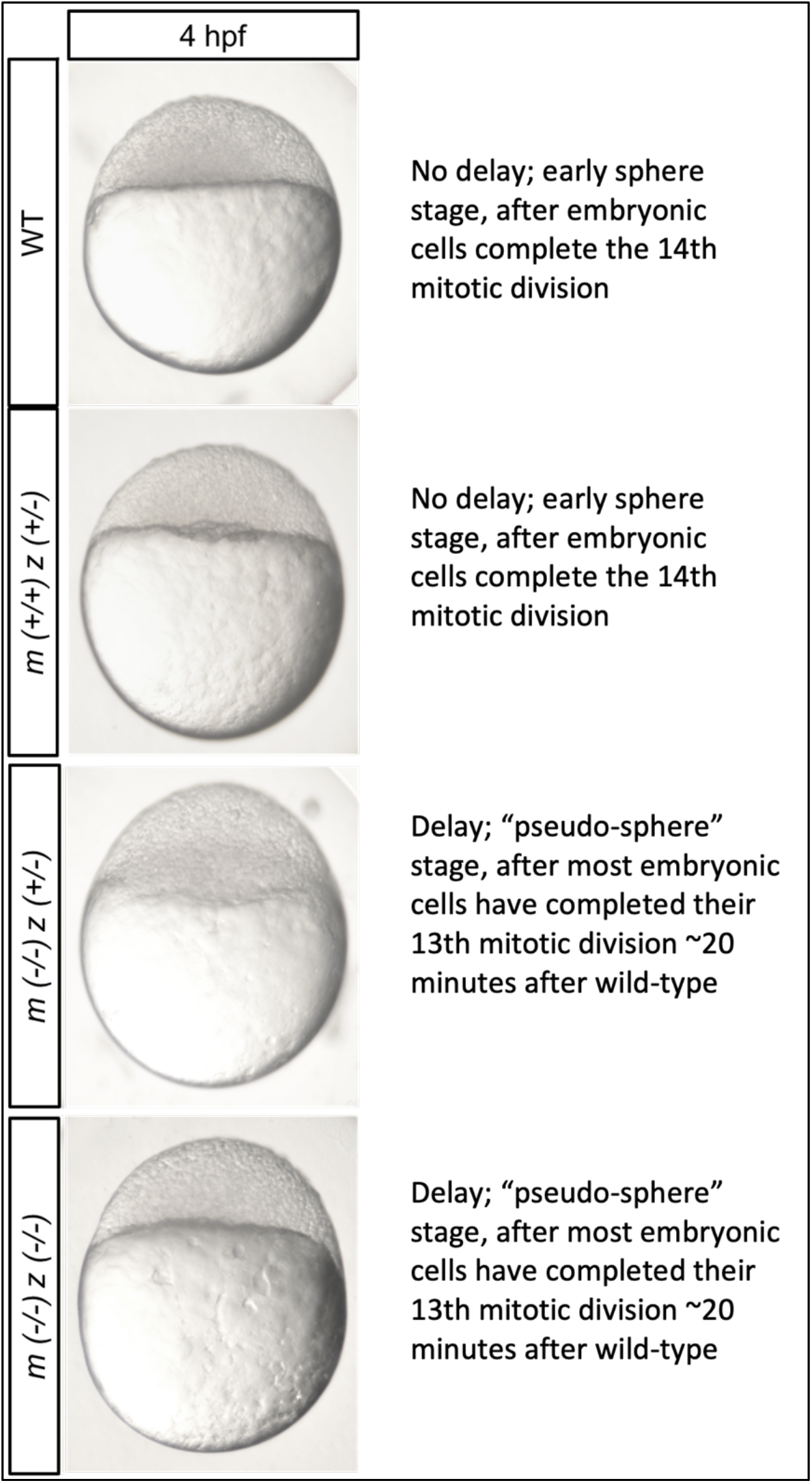
Observing developmental stages and cell characteristics during zebrafish blastula development by bright field microscopy. Micrographs previously presented in this work and in Zhao *et al*. 2017^*2*^. Direct comparison of embryos at 4 hours post-fertilization (at 29 +/- 1.5°C), illustrating wild-type and maternal (+/+) zygotic (+/-) *ythdf2(ch200)* embryos at sphere stage (top two images) versus maternal (-/-) zygotic (+/-) and maternal (-/-) zygotic (-/-) *ythdf2(ch200)* embryos at “pseudo-oblong” stage (bottom two images). Together, this comparison provides a proper control for genetic background.

Moreover, previously unpublished data presented in **Figure 4** counters the *in situ* hybridization results of Kontur *et al*. (Fig. 2E in that work). We observe specific, aberrant gene expression patterns for developmentally important genes (*chrd, aplnra*, and *foxa3)* in maternal *ythdf2(ch200)* mutant embryos versus wild-type embryos at the same developmental stages (*not* at the same number of minutes elapsed since fertilization). Mutant gene expression patterns gene are not the same as those observed in younger embryos (https://zfin.org/), and they cannot be readily explained by invoking a natural developmental timing delay that arises in the hybrid progeny of two different wild-type zebrafish fish strains. Additionally, we have observed RT-PCR-based RNA misprocessing events in maternal *ch200* embryos (data not shown), which are also not easily explained by invoking strain background effects. In summary, these additional data provide further evidence for aberrant developmental progression in embryos from *ythdf2(ch200)* homozygous mutant females.

**Figure 4.**
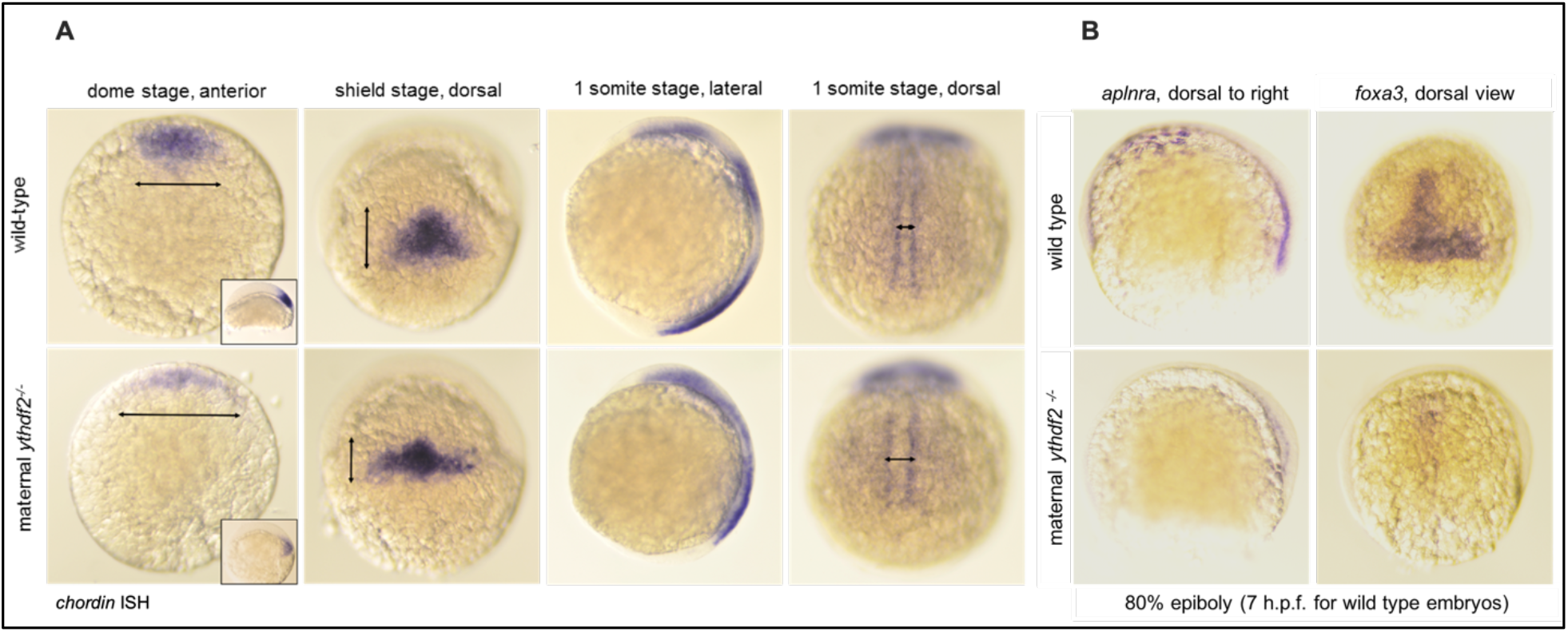
Aberrant gene expression in ythdf2(ch200) m(-/-) z(+/-) embryos crossed into the *AB background versus *AB wild-type controls is revealed by in situ hybridization for developmentally important genes. (A) *chordin (chrd)* RNA probe, previously described; (B) *aplnra* RNA probe, created for this study, and *foxa3* RNA probe, previously described. Comparisons are between staged-matched, not time-matched, wild-type and mutant embryos spanning late blastula to mid-gastrulation time points.

### Zygotically transcribed m^6^A-modified genes, including “maternal” genes that are resynthesized zygotically, must be accounted for in pre-versus post-ZGA time point comparisons

It is impossible to make direct inferences about the fates of maternally m^6^A-modified RNAs by comparing across pre- and post-ZGA time points without accounting for the synthesis of m^6^A-modified zygotically transcribed genes. This includes “maternal” genes that are re-synthesized zygotically at ZGA, which may or may not differ in m^6^A composition from their maternally deposited counterparts.

Comparisons of RNA populations pre-versus post-ZGA (*i.e*., zebrafish 2 hpf vs. 4 hpf or 6 hpf) were performed often in Kontur *et al*. (Fig. 1A-C, Fig. 2B and D, Fig. 3B, Fig. 5B, Fig. 6C-F, and Supplemental Figures). However, nascent zygotic transcription and continued clearance of maternal RNAs occur simultaneously across this time interval. Maternal RNAs (both un-modified and m^6^A-modified) are found *both* pre- and post-ZGA, and new, zygotically transcribed genes, which may or may not have also been deposited maternally, are themselves either m6A-modified or un-modified. Hence, even selecting specifically for m6A-containing RNAs cannot reveal the dynamics of m6A-modified transcripts during this complicated developmental time. Moreover, the metabolic mechanisms affecting m^6^A-containing RNAs may very well differ at least in part before versus after ZGA^5–7^. Blocking transcription at ZGA is a logically sound experimental strategy in this context, but it is also a radical manipulation that affects maternal RNA degradation and m^6^A metabolism by itself, as zygotically-derived factors are needed to further the maternal RNA degradation program^8^, and *ythdf2* increases in expression just after ZGA in zebrafish embryos^1^, presumably to further regulate m^6^A-containing RNAs.

To concretize this point with data collected in Zhao *et al*. from wild-type embryos, we found that of 4,762 genes with transcripts detected throughout the interval from 0 to 8 hpf, 1,806 (37.9%) are not m^6^A-modified either before or after ZGA, 1,675 (35.2%) are m^6^A-modified continuously from 0 to 8 hpf, 476 (10.0%) are only found m^6^A-modified maternally (0-2 hpf), and 805 (16.9%) are only found m^6^A-modified zygotically (6-8 hpf). Thus, 56.6% of maternally deposited transcripts that contain m^6^A remain m^6^A-modified from fertilization to well past ZGA (without respect to whether they are re-synthesized [with m^6^A] at ZGA). Importantly, however, an appreciable fraction of transcripts, *a)* are maternally deposited as m^6^A-modified and then retranscribed after ZGA without m^6^A (16.1%), or instead, *b)* are maternally deposited un-modified, but upon ZGA are re-transcribed to contain m^6^A (27.2%). These data highlight a markedly dynamic population of m^6^A-containing mRNAs across the period of ZGA during zebrafish development, which must be accounted for experimentally in order to understand the source and fate of m6A-contaning RNAs in the early embryo.

### Poly(A)-based RNA sequencing is problematic in the early embryo

Dynamic poly(A) tail length changes in the early embryo, including deadenylation decoupled from RNA decay and the unusual phenomenon of cytoplasmic polyadenylation, make poly(A)-based sequencing particularly unsuitable for answering general questions about RNA metabolism during the earliest stages of development. Most maternal mRNAs are initially inherited with unusually short poly(A) tails. These tails then undergo dramatic remodeling via lengthening or shortening during early embryogenesis, both before and after zygotic genome activation (ZGA), to govern their translational prowess (increased for messages with long poly(A) tails), ability to remain in a stably “repressed” state for future translation (typical of short-intermediate poly(A) tails), or degradation (typical of very short poly(A) tails)^9,10^. After ZGA, the embryo adds new, zygotically-transcribed mRNAs to the remaining maternal mRNA pool, and, among other changes, poly(A) tail shortening is found to promote active transcript destabilization.

In light of these complicated poly(A) tail dynamics, heavy reliance on poly(A)-selected mRNAs at early developmental stages during scientific work is problematic. In Zhao *et al*., we used only ribo-minus-based RNA selection followed by deep sequencing to address the dynamics of maternally deposited RNA. Indeed, we find that even in Kontur *et al*., which employed both poly(A)-based and ribo-minus RNA selection, stronger differences in global mRNA abundance are observed between experimental and control samples using ribo-minus based, rather than poly(A)-based, RNA selection (see especially their Supplement).

A particular example of the limitations of poly(A)-based RNA sequencing in early development can be discerned in Kontur *et al*. Figures S4. In Fig. S4A, poly(A)-based RNA selection at pre-ZGA time points initially shows a steady increase in the levels of *ythdf1, ythdf2*, and *ythdf3* mRNAs from 0-~3 hpf. However, there can be no actual increase in mRNA levels for these genes during these time points, because there is (essentially) no RNA synthesis yet in the embryo^11^. Instead, the ribo-minus-based RNA selection in Fig. S4B captures the RNA levels of the different transcripts more accurately, revealing a basically static amount of *ythdf1, ythdf2*, and *ythdf3* messages from fertilization until ZGA, at which point there is a dramatic increase in the levels of *ythdf2* (but not *ythdf1* or *ythdf3*).

### Results from derived reporter constructs should be interpreted with caution

We note that the m^6^A-containing RNA reporter construct used in Kontur *et al*. (Fig. 1E and H, Fig. 2G in that work) is dissimilar in m^6^A distribution from typical endogenous m^6^A-modified transcripts. Multiple studies have found that *in vivo*, roughly 50-70% of m^6^A-modified residues are located at the 3’ end of the coding region and within the first third of the 3’ UTR, roughly 20-40% are found throughout the coding region, and 10-15% are located at the transcription start site (TSS) and within the 5’UTR^12,13^. As has been increasingly documented, distinct locations of m^6^A in different parts of the transcript affect different aspects of mRNA metabolism, sometimes in contradictory ways^14,15^. The specific density of m^6^A sites seems to tune the proper binding of YTHDF and other RNA binding proteins^16,17^ to influence transcript decay^18,19^ and/or translation^20–23^ and/or RNA secondary structure^24,25^.

In contrast to the usual m^6^A distribution, the m^6^A reporter used in Kontur *et al*. contained no m^6^A at either the TSS, in the 5’UTR, or within the first ~75% of the reporter coding sequence (with the exception of the start codon). Although the reporter does possess an m^6^A bias at the 3’ transcript end, every adenosine within the reporter’s 3’UTR was able to be m^6^A modified (depending on the stochastic incorporation of A versus m^6^A during *in vitro* transcription), preventing m^6^A skew towards the beginning of the 3’UTR. Both the artificial number and distribution of m^6^A-containing sites in this reporter would create significant risk that it cannot be regulated as an endogenous m^6^A-containing transcript would be.

### High-throughput sequencing assays should be performed at least in duplicate

By examining Table S2 of Kontur *et al*., we find that of the 18 high-throughput sequencing experiments involving *ythdf2(ch200)*, 16 lacked any replication. Only two experiments seem to have been run in duplicate: ribo-minus “background-matched wild-type” at 6 hpf and ribo-minus “MZythdf2 (*ch200)’’* at 6 hpf. All other deep sequencing experiments involving *ch200* or “background-matched” controls, using either ribo-minus RNA or poly(A)-selected RNA at any developmental stage, are catalogued to have only been conducted a single time.

## Conclusions

Our intention in Zhao *et al*. was to bring to light a new molecular actor in maternal RNA degradation, Ythdf2, which works in part prior to zygotic genome activation to clear maternal RNAs, and allows, directly or indirectly, embryos to undergo proper zygotic genome activation at the proper time. We understood then that Ythdf2 “is not the sole regulator of methylated mRNA stability,” and we did not claim that Ythdf2 is “mandatory for clearance of all methylated maternal transcripts” or that “Ythdf2 alone is critical to developmental timing,” as was suggested by Kontur *et al*. Indeed, we have always agreed with Kontur *et al*. in believing that the roles of Ythdf proteins, and other m^6^A “readers” known and yet to be discovered, “must be fully defined to completely elucidate how methylation governs mRNA clearance during key cellular transitions.”

One notable development since our original paper was published in 2017 is the finding that terminal uridylation is responsible for a large fraction of maternal RNA destruction, especially after ZGA^26,27^. Indeed, Tut4/7-mediated uridylation affects the largest proportion of maternal transcripts yet known compared to other regulatory mechanisms. In addition, we note that work published in the mouse model at nearly the same time as ours corroborates YTHDF2’s key roles in regulating transcriptome turnover for oocyte maturation and early zygotic development^28^.

## Notes

### Competing Interest Statement

The authors have declared no competing interest.

